# CRE mice exhibit hyperactive and impulsive behavior affecting their learning and retention performances

**DOI:** 10.1101/2024.02.27.582228

**Authors:** Frédéric Desor, Aseel El Hajj, Ameziane Herzine, Fathia Djelti, Vincent Bombail, Isabelle Denis, Thierry Oster, Catherine Malaplate, Marie-Claire Lanhers, Frances T. Yen, Thomas Claudepierre

## Abstract

CRE recombinase is a protein that recognizes and mediates site-specific recombination between loxP site sequences. The *Cre/loxP* recombination system has become a useful tool for genetic manipulation. Spatial regulation of recombination can be achieved by using cell type-specific promoters that drive expression of CRE in the tissue of interest. The temporal regulation can be obtained with CreER recombinase, which consists of Cre fused to mutated hormone-binding domain of the estrogen receptor (ER). In the more improved versions of the construct, the CRE*-*mediated gene regulation can be controlled both spatially and temporally, by combining tissue-specific expression of a CreER recombinase with its tamoxifen-dependent activity. We recently generated and characterized an astrocyte specific mutant of the lipolysis-stimulated lipoprotein receptor *lsr* gene by crossing Glast ERT2 mice with floxed *lsr* mice (El Hajj et al., 2022). During the behavioral analysis of generated mice, we identified specific hyperactive traits in the Glast ERT2 mice (CRE mice) that prevented them from being used as a control group. Here we further assessed the hyperactive trait of those CRE mice using a battery of behavioral tests. We showed that CRE mice exhibited hyperactive behavior combined with attention-deficit, sleep disturbance and impulsivity that affect their learning and memorization performances. These mice may therefore serve as a model to study attention deficit / hyperactivity disorder. Our work also pointed out the need for proper behavioral analysis of control groups in transgenic animal generation to avoid misinterpretation and misattribution of behavioral traits.

## 1. Introduction

Vectors expressing CRE recombinase have been utilized to turn gene expression on or off. The CRE recombinase is a protein that recognizes and mediates site-specific recombination between *loxP* site sequences in bacteriophage P1 [1]. CRE-mediated recombination between two *loxP* sites can result in gene deletion, insertion, translocation, and inversion depending on the location and orientation of the loxP sites. The *Cre/loxP* recombination system has turned into a useful tool for directed genetic manipulation *in vitro* [2] and *in vivo* [3]. The *Cre/loxP* site-specific recombination system is particularly useful for conditional somatic mutation in mice. Spatial regulation of recombination can be achieved by using cell type-specific promoters that drive expression of CRE in the tissue of interest, such as using the *Glast* promoter to drive CRE expression in astroglial cells that specifically express this glutamate transporter [4]. Temporal regulation can be obtained with CreER recombinase, which consists of Cre fused to the mutated hormone-binding domain of the estrogen receptor (ER). The CreER recombinase is inactive but can be activated by the synthetic estrogen receptor ligand tamoxifen (TAM). The more improved versions of the chimeric CRE recombinase have been developed, including CreER^T2^ [5]. By combining tissue-specific expression of a CreER recombinase with its TAM-dependent activity, the CRE-mediated gene regulation can be controlled both spatially and temporally [6]. For instance, induction of *GLASTCreER*^*T2*^ mice with TAM drives, in a temporally-dependent manner, the glia-specific expression of CRE enzyme [4].

At the time we established the behavioral phenotyping of a new glia-specific conditional knockout mouse (cKO) using the *GLASTCreER*^*T2*^construct [7], we chose to include CRE littermates as a control group in the behavioral studies. The only difference between them and the group of interest (cKO) was, supposedly, the absence of deletion of the target gene in the CRE group Despite this, CRE mice exhibited abnormal behavior and low results in all tests requiring attention. They showed signs of hyperactivity that prevented us from using them as a proper control group. However, this hyperactive trait could turn the CRE mice into an interesting model to study attention deficit / hyperactivity disorder (ADHD). This disorder is characterized by a persistent pattern of inattention and/or hyperactivity-impulsivity that has a direct negative impact on academic, occupational, or social functioning [8]. In children, attention deficit may result in poor school performance. In addition, ADHD is associated with mental disorders and substance abuse in adulthood [9]. Since CRE mice exhibited excessive motor activity that could account for hyperactive trait and performed poorly in attention-requiring tasks that could reflect distraction and inattention, we performed an extensive behavioral analysis of the CRE mice by comparing their score to control wild type in specific attention- and activity-related tests. Results obtained confirmed their hyperactive behavior, specific memory and learning deficits, but ruled out anxiety trait or sensory deficits. Altogether the profile of CRE mice resembled ADHD, highlighting their potential usefulness as model of this cognitive disorder, in addition to existing animal models [10]. It also pointed out possible bias introduced in behavioral studies where abnormal behavior in generated cKO was linked to CRE and not to the excision of the target gene.

## 2. Materials and Methods

### 2.1. Animals

Initial C57Bl6J genitors of Glia-specific CRE/+ transgenic mice (Tg(Slc1a3-cre/ERT2)45-72Fwp) [4] were a kind gift from Dr FW Pfrieger (INCI, Strasbourg). C57Bl6J WT control mice of the same genetic background were purchased from Charles River Laboratories (Saint Germain Nuelles, France). Male CRE and WT mice were housed in certified animal facilities (#B54-547-24), under controled environmental conditions including a constant temperature (22 ± 2 °C), and a relative humidity of 50 ± 5 % with *ad libitum* access to food (standard chow diet, Envigo Teklad, Gannat, France) and water. Animals were maintained on a standard 12-h light/dark cycle: dark cycle starting at 12:00 P.M. (noon) and ending at 12:00 A.M. (midnight). Animal studies were conducted in accordance with the European Communities Council Directive (EU 2010/63) for the use and care of laboratory animals. All experimental procedures were approved by the institutional ethical board (authorization number APAFIS #12079-201711081110404).

### 2.2. Tamoxifen (TAM) injection

TAM (Sigma, St Louis, MO) was dissolved in a 9:1 (v/v) sunflower oil (Sigma, St Louis, MO) and ethanol (Carlo Erba Reagents, Val de Reuil, France) mixture at a concentration of 15 mg/mL at 37 °C, sterile-filtered (0.22 μm), and stored at 4 °C for up to 7 days in the dark. A 23G needle tuberculin syringe (Henke Sass Wolf, Tuttlingen, Germany) was used for intraperitoneal injections. At the age of 8 weeks, all mice were injected for 5 consecutive days (every 24 h) with 150 μg of TAM per g of body weight [4]. Each mouse was randomly chosen from a different litter to avoid any litter-specific bias (litter effects).

### 2.3. Behavioral tests

Two weeks after TAM induction, behavioral tests detailled in Table 1 were performed over an 8–month period using the same mice during the entire study (15 WT and 14 CRE mice). All behavioral tests were performed 1 h after the beginning of the dark cycle using only red light.

**Table 1.** Timeline of behavioral tests.

| Behavioral test | Age of animals |
| --- | --- |
| Open field test (OFT) | 3 months |
| Y-maze | 4 months |
| Object recognition test | 4.5 months |
| 3-chamber sociability and social novelty test | 5 months |
| Home cage activity | 6 months |
| Barnes maze | 11 months |

#### 2.3.1. Open field

The general locomotor activity was assessed on 3-month old animals (15 WT and 14 CRE mice) in a white circular open field (80 cm diameter × 50 cm high walls) coupled with a Smart video tracking system (Bioseb, Vitrolles, France). Using this software, the radius of the circular area was virtually divided into three sections to establish three circular areas: peripheral (Z1), median (Z2) and central areas (Z3). Four adjustable lamps, arranged on the ceiling allowed indirect illumination of the area. Luminous intensities were 20, 35 and 80 lux in Z1, Z2 and Z3 respectively. Testing was conducted between 2:00 P.M. and 5:00 P.M. Animals were placed in the peripheral zone (Z1) of the device, head facing the wall, and ambulation was monitored during 3 min. Parameters recorded for the locomotor activity assessment were: total traveled distance, cumulated duration in each area, as well as latency of the first entry and number of entrances in Z3. A moving fast threshold was established for speed above 14 cm/s, allowing the measure of the total time moving at high speed.

#### 2.3.2. Y maze

Immediate spatial working memory performance was evaluated on 4-month old animals (15 WT and 14 CRE mice) using a classic Y maze. The device was composed of three opaque black plexiglass arms (40 L x 9 W x 16 H cm, and positioned at equal angles) with a floor made of transparent plexiglass, allowing the animal to see geometric patterns depending on the arm, as proximal clues. The Y-maze task was conducted between 2:00 P.M. and 5:00 P.M., in a dimly illuminated room (80 lux in maze arms). Mice were placed at the extremity of one arm, head facing the wall of the maze, and allowed to move freely through the apparatus during a 5-min session. The percentage of spontaneous alternation was calculated as the ratio of successful overlapping alternations by the total possible triplets (defined as the total number of arm entries minus 2) multiplied by 100. An arm entry was considered to be completed when the four paws of the mouse were all placed in the arm.

#### 2.3.3. Object recognition test

The object recognition test was performed on 4.5-month old animals (15 WT and 14 CRE mice) according to the procedure of Leger *et al*. (2013) [11]. Briefly, each mouse was first habituated to the device (acclimation) before being subjected to an acquisition phase during which two similar objects were presented to it, then an hour later, had to discriminate a new object from the one already acquired (phase of recognition memory). The procedure was conducted from 1:00 P.M. to 5:00 P.M. in an empty squared opaque plastic box (30 L x 30 W x 26 H cm) placed in a dimly illuminated room (25 lux in the center of the arena). During the acclimation session, the arena was left empty and the mouse was allowed to freely investigate its new environment for 3 min in order to avoid any later potential neophobic response. Monitoring of the mice placement was carried out using the video tracking system to highlight, if existing, any initial place preference in the device especially in the 2 opposite corners of the arena (namely position A and B) where objects would be placed in the following steps of the test. Fifteen minutes after the acclimation phase, an acquisition session was conducted in which the mouse was presented two similar objects (either two Lego blocks or two litter-filled tissue culture flasks, 15 cm high to prevent the mouse from climbing on it) positioned in positions A and B. The animal was allowed to explore the objects for 3 min. During this session, the monitoring of both similar objects was performed to enlighten any initial object preference that would induce an experimental bias in the later recognition memory test. One hour later, one of the known objects was randomly removed and a new one (either Lego or Falcon) was placed at the same position. The novel object recognition session was then performed, in which exploratory behaviors toward objects were recorded via the Smart video tracking system. To this end, we used the TriWise option, which allowed tracking the position of the mouse’s head relatively to the animal’s center. According to Leger *et al*., the mouse was considered as exploring the object when its head was directed toward the object at a distance less than or equal to 2 cm. Mice memory was assessed by calculating the ratio of “time spend exploring the novel object” to “total time of objects exploration” (recognition index) obtained during the discrimination task. We also focused on mice interest toward their environment during both acquisition and discrimination phases by adding times of exploration toward both objects, with no interest for a possible object preference.

#### 2.3.4. Social interaction / memory test

The social memory assessment was performed on 5-month old animals (15 WT and 14 CRE mice) using a device made of an open-top transparent plastic box (63 L × 42 W × 22 H cm), divided into three successive chambers by two opaque partition walls (Imetronic, Reganeau, France). Small openings (5 x 5 cm) in these partitions allowed mice to access each compartment of the apparatus. Both opposite chambers were equipped with a cylindrical wire cup made of chrome bars spaced 1 cm apart (11 cm H; 11 cm bottom diameter). We followed the Kaidanovich-Beilin’s procedure [12], consisting of three phases, *i*.*e*., a first phase of familiarization to the device, a phase of sociability assessment and a phase of social memory testing. During the familiarization phase, the mouse was placed in the middle chamber and was allowed to freely explore the whole device for 5 min to reduce anxiogenic response to novelty. Again, we used video tracking to highlight, if existing, a place preference (cumulated time of presence in each of the opposite chambers) which could potentially induce bias in the socialization and/or social memory results. Following this familiarization phase step, the mouse was replaced in its home cage for 5 min before sociability assessment phase. Once again, the tested animal was placed in the middle chamber of the apparatus, the device now containing an unknown congener previously positioned under one of the wire cups. The TriWise option from Smart video tracking system software was used to record the sniffing time around both wire cups in a range of 2 cm and for a 5 min duration. Thus, the social motivation was assessed via the time spent sniffing the cup containing congener versus the empty cup. At the end of the socialization phase, the mouse was replaced in its home cage for one hour before the social discrimination test was performed. In this aim, the now-known congener was placed under the same wire cup used in the socialization step, while an unknown congener was placed in the second cup of the opposite chamber. Again, the tested animal was placed in the device’s central chamber and allowed to explore the device for 5 min during which sniffing time around the wire cups (2 cm) was recorded and considered as exploration time. Total investigation time and the ratio of social discrimination (time spent sniffing unfamiliar mice relative to total investigation time) were calculated.

#### 2.3.5. Spontaneous home cage activity

The assessment of the spontaneous home-cage activity was performed on 6-month old animals (15 WT and 14 CRE mice) using four beam-break activity monitors (Promethion, Sable Systems International, North Las Vegas, USA). The array of each monitor was composed of three independent infrared (wavelength 900 nm) beams (X + Y + Z axes), whose captors were spaced 0.25 cm on each axis. The recording sessions took place in the animal room to avoid any disruptive or stress effect that could occur if moving the mice to another room. At 11:00 A.M., cages were placed in the center of each beam-break monitor. The duration of monitoring was set at 24 h, beginning at 12:00 A.M. (light “off”). The subsampling factor was set at 1 s and raw data were processed using ExpeData software (Sable Systems International). According to the manufacturer’s instructions, ambulation exceeding 1 cm/s was considered as directed locomotion (ped-meters), while values below this threshold were considered as non-ambulatory movements, *i*.*e*., fine activity (grooming, scratching or feeding). The variable “all meters” was established as the sum of all distances traveled within the beam-break system including fine movement as well as direct locomotion. Moreover, using guidelines from Pack *et al*. (2007) [13], any episode of continuous inactivity for ≥ 40 s was considered as sleep and expressed as percentage of the cycle. Given the wide difference of activity in mice between the light and dark phases, data were initally examined according to each phase of the cycle, then to the total circadian cycle. Also, in order to allow genotype comparisons for each activity-related variable, the 12 values within a cycle’s phase were summed up. Thus, the level of significance of the subsequent unpaired *t*-test (CRE vs WT) was equal to the main factor “genotype” from the two-way ANOVA with repeated measures of the activity-related variable, and thus, not mentioned.

#### 2.3.6. Barnes maze

Eleven-month old animals (15 WT and 14 CRE mice) were tested on a modified Barnes maze, made according to the work of Youn *et al*. (2012) [14]. The maze consists of a white circular area (radius of 56 cm), positioned in height (40 cm), and pierced with 44 openings (5 cm diameter) placed according to 3 circumferences (peripheral, median and central), and following a similar pattern for its four quadrants. Only one escape hole was provided with a small mesh staircase leading to the escape chamber, a dark and restricted space in which the mice were left for one minute after having found the target hole. According to Youn’s procedure, the escape hole was never placed on the peripheral circumference to avoid experimental bias induced by serial thigmotaxic explorations. The apparatus was mildly illuminated (100 lux at the center), and 3 rotating fans placed equilaterally around the device induced aversive conditions to obtain sufficient motivation in the mice to seek for the escape hole during the trial sessions. Four distant clues (black geometric shapes on white panels) were placed on the walls around the device at the cardinal points, 30 cm higher from its surface to allow mice’s allocentric navigation. The escape hole was set under one of the clues and its location never changed during all the test sessions. The mouse was placed on the modified Barnes maze device inside a small opaque plastic cylinder (10 cm diameter / height) to avoid any pre-trials spatial recognition. The location of the trial’s starting point was set at the center of the quadrant opposite to the target quadrant. The test started with the withdrawal of the cylinder, which was performed when the mouse had its head placed in the opposite direction to the exit hole. In this way, the animal was required to perform an initial half-turn to move to the escape hole. Mice were given 3 min to find the escape hole. When an animal did not succeed in the imparted time, it was gently positioned and maintained by the tail for 10 s in front of the escape hole, *i*.*e*., facing its specific distant clue, before being allowed to enter the hole. Familiarization and acquisition sessions took place for three consecutive days (D1, D2 & D3), in which mice were given three trials a day to acquire the location of the escape hole. Time to find the escape hole were recorded for each trial. Two probe tests were performed on the fifth (D5) and eighth days (D8) consisting of only one trial/day to evaluate memory retention. The latter is expressed as the percentage of the time spent in the target quadrant compared to untargeted quadrants according to the total time needed for the trial.

### 2.4. Statistical analyses

All variables were tested for its distribution by the Kolmogorov-Smirnov test. Since all variables followed a normal distribution, parametric statistics were used for the analyses of this work and results are presented as mean ± SEM. Data obtained from the spontaneous home cage activity and Barnes maze acquisition sessions were analyzed using the two-way repeated measures analysis of variance, considering genotype and time as main factors. Intra-group performances in the Barnes maze were analyzed using a 1-way ANOVA with repeated measure, with time as unique main factor. If significant, a post-hoc Tuckey test was used to enlighten significant differences of the variable between specific times. Data resulting from non-time-dependent comparisons were performed via unpaired Student *t* test for strain comparisons, while intra-group performances were achieved using paired Student *t* test. Statistics were performed using StatView software (version 5.01) and in all analyses, *p* < 0.05 was considered statistically significant.

## 3. Results

### 3.1. Open field

When comparing the total distances traveled in the open field during the 3 min of the test, CRE mice traveled a greater distance when compared to controls (*p* = 0.027, **Table 2**). By examining the distribution of the distances traveled according to the area of the device, it appeared that compared to controls, CRE mice moved significantly more in the peripheral Z1 area (*p* < 0.0029) and less in the central Z3 area (*p* = 0.0074). However, no difference was found regarding the distance travelled in the middle area (Z2) of the open field (*p* = 0.76). This thigmotaxic behavior was still found when examining the times of presence in each zone according to the mouse strain. CRE mice also exhibited a significantly longer time in Z1 (*p* = 0.0031), with shorter times in Z2 (*p* = 0.06) and Z3 (*p* = 0.0016) when compared to controls (Table 1). The analysis of the two variables “latency of the first entry”, as well as “total number of entries” in the central zone, as representative of the level of anxiety in rodents, revealed no difference between strains (latency of the first entrance in Z3, *p* = 0.982; total number of entries in Z3 during the 3-min test, *p* = 0.095). Finally, the analysis of the total time spent by animals moving at high speed, i.e., > 14 cm/s threshold, was found to be significantly different (*p* = 0.0038), with the CRE animals spending a total of 57 s at high speed while control animals exhibited a duration of 33 s. This most likely explains the difference observed between the two groups concerning the distance travelled in the open field.

**Table 2.**
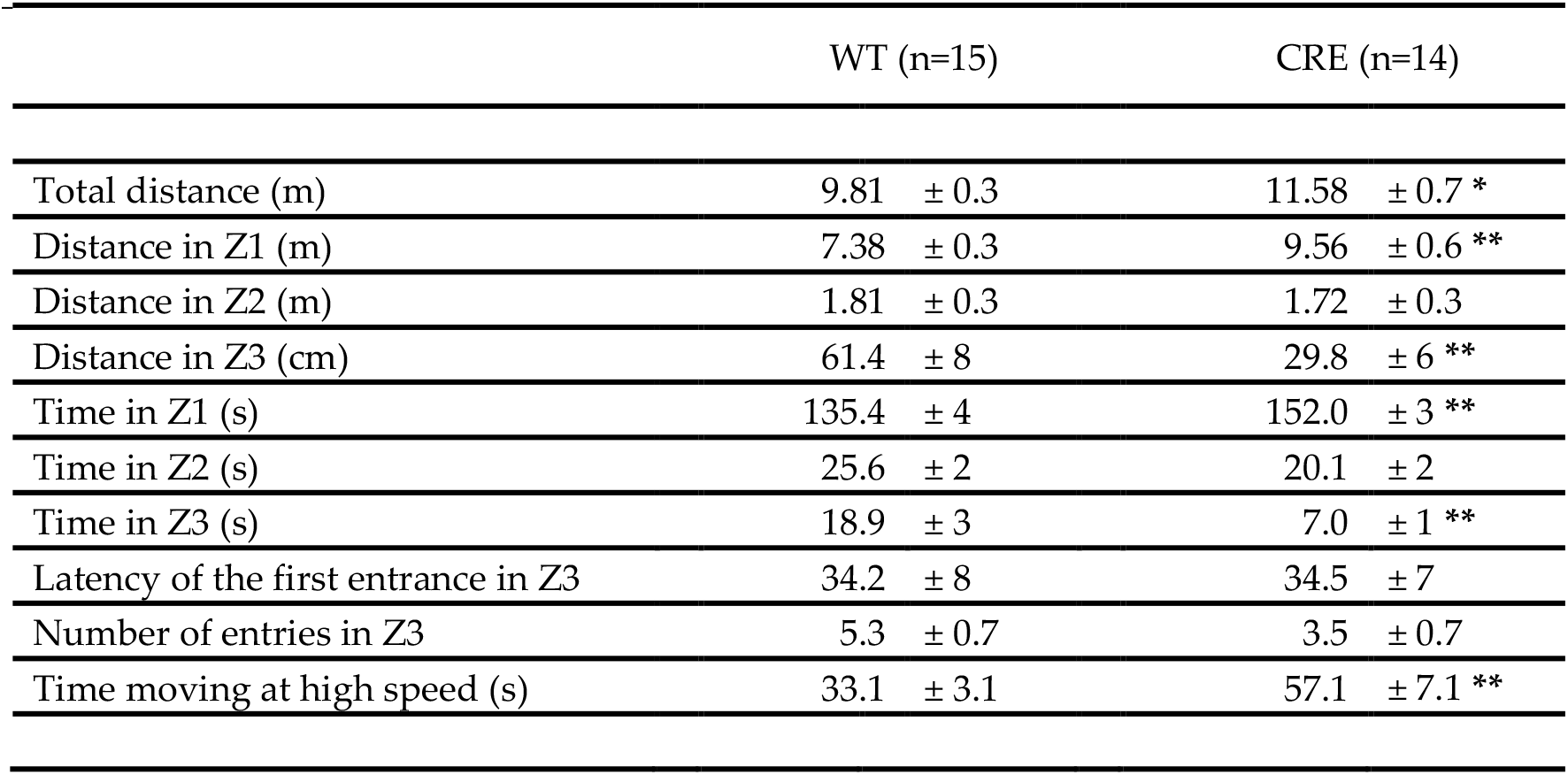
Behavior of CRE and WT in open-field test (CRE: *n* = 14; WT: n = 15). Data are reported as mean ± SEM. \**p* < 0.05, \*\**p* < 0.01, significantly different from control group (unpaired *t* test).

The total number of triplets performed by mice in the Y-maze device revealed a higher ambulatory activity (*p* < 0.0001), where CRE mice completed up to 26.3 triplets (± 1.9) and WT control mice up to 18 triplets (± 1.1) (**Figure 1A**). Nevertheless, this performance of CRE mice was also accompanied by a poor immediate spatial memory, since the percentage of spontaneous alternation was found close to the chance threshold for CRE mice (X=53.7 ± 2.6 %), while WT mice exhibited higher performances of 67.2 ± 2.5 %, *p* = 0.0008, **Figure 1B**).

**Figure 1.**
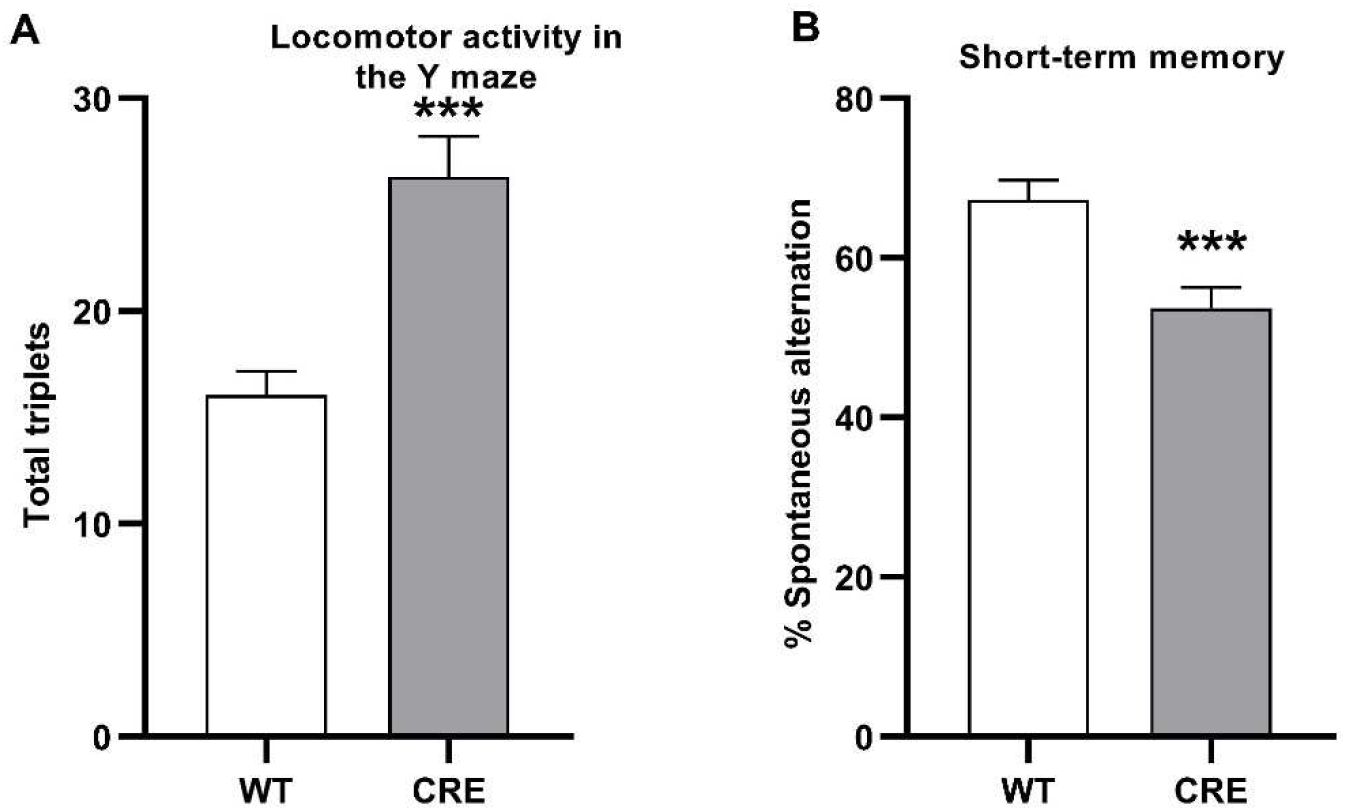
Effect of *Cre* genotype compared to WT on the behavior in the Y maze (CRE n=14, WT n=15). Data are reported as mean ± SEM. \*\*\**p* < 0.001 significantly different from control (Fisher *post hoc* test).

### 3.3. Object recognition test

No place preference in the device was observed during the acclimation step, neither in WT control mice (*p* = 0.17) nor in CRE mice (*p* = 0.23). Also, no object preference was revealed during the acquisition session neither in control mice (*p* = 0.30) nor in CRE mice (*p* = 0.15). Concerning their interest towards their environment, control mice showed a longer exploration time (X = 40.7 ± 5 s) when compared to CRE mice (X=24.8 s ± 3 s, *p* = 0.019, **Figure 2A**). A difference was still found during the discrimination task, but did not reach the level of significance (*p* = 0.10, **Figure 2B**). The discrimination task, performed 1 h after the acquisition phase, revealed that control mice spent 63.5 ± 3 % of their exploration time towards the novel object, while CRE mice recognition index was found close to the chance threshold (50.7 ± 5 %, *p* = 0.04, **Figure 2C**).

**Figure 2.**
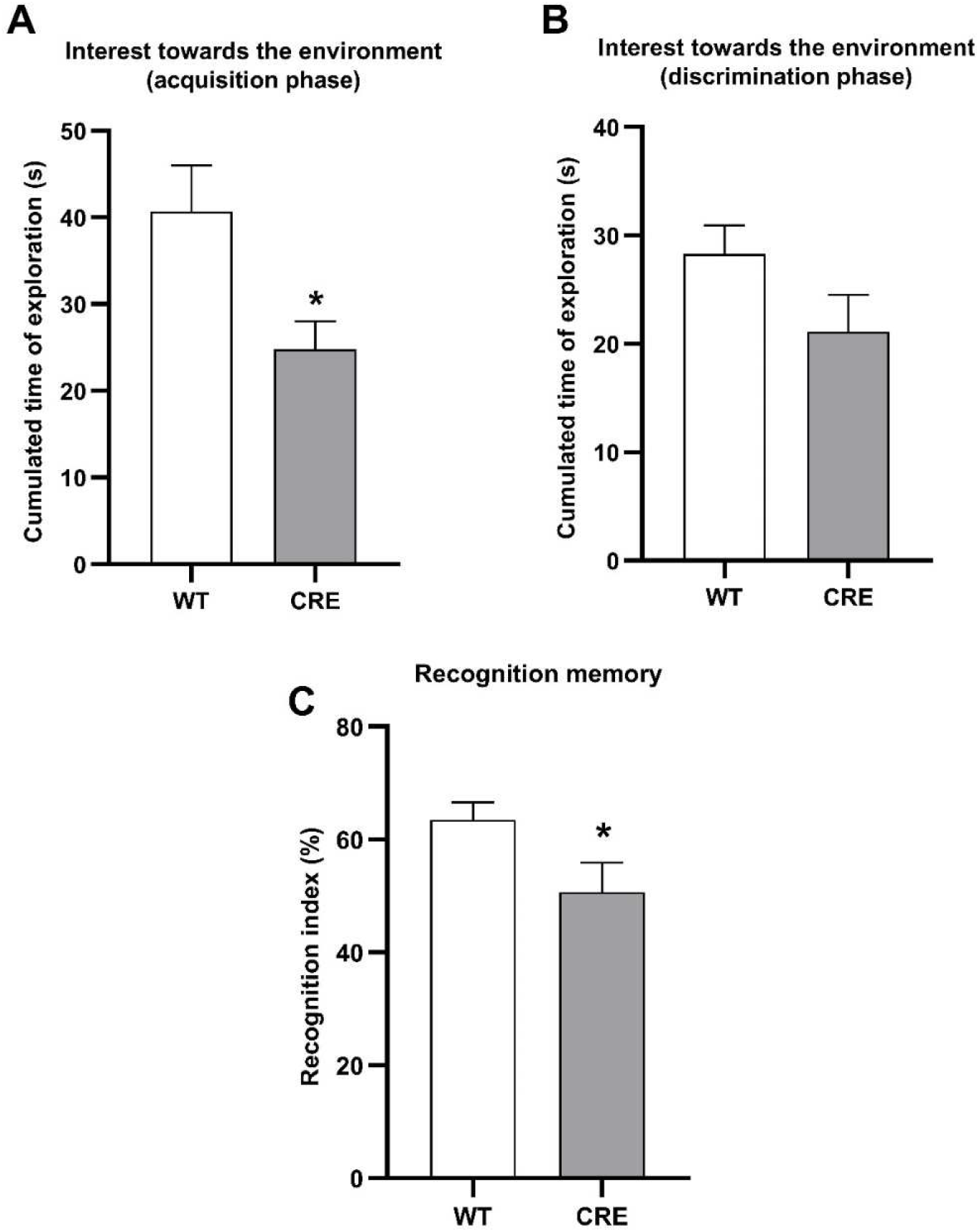
Effect of *Cre* genotype compared to WT on the interest towards the environment (A & B) and recognition memory (C) in the object recognition test (CRE n=14, WT n=15). Data are reported as mean ± SEM. \**p* < 0.05 significantly different from control (Fisher *post hoc* test)

### 3.4. Social discrimination test

Since CRE mice exhibited a lack of interest toward the inanimate environment of the object recognition test, we performed a two-trial social memory test to assess the possible influence of the type of stimuli on CRE mice attention (**Figure 3**).

**Figure 3.**
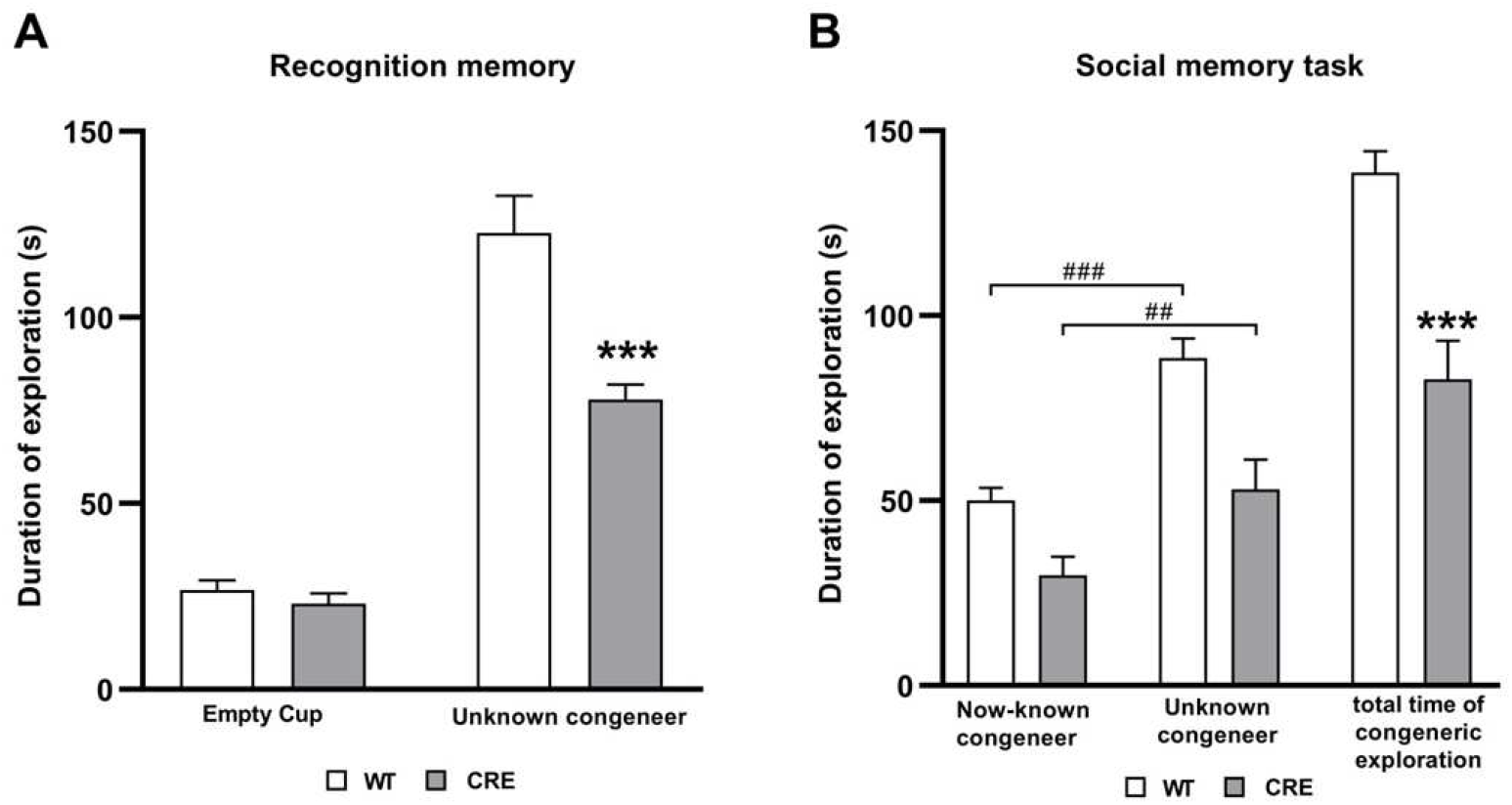
Effect of *Cre* genotype compared to WT on times of congener exploration during sociabilisation and discrimination phases (CRE n=14, WT n=15). Data are reported as mean ± SEM. \*\*\**p* < 0.001 significantly different from control (Fisher *post-hoc* test). ^##^*p* < 0.01; ^###^*p* < 0.0001, intra-group significant difference (paired t-test).

Initial comparison of mice movments in the empty apparatus during the familiarization phase did not reveal any place preference for a specific compartment (data not shown). Evaluation of mice sociability showed that CRE mice spent more time exploring the wire cup containing an unknown congener than the empty one (*p* < 0.0001). This same discriminative behavior was also found in controls (*p* < 0.0001). Nevertheless, it was observed that the CRE mice spent significantly less time (*p* = 0.0004) to explore the congener (78 ± 4 s) than the controls (123 ± 10 s). However, no difference was found between the groups with regard to the time spent in exploring the empty wire cup (*p* = 0.33, **Figure 3A**). Social memory task revealed that despite a lower time of congener exploration during the social motivation assessment, CRE mice were nonetheless able to discriminate the congeners (*p* = 0.0038). Not surprisingly, this result was also found in WT mice (*p* = 0.0004, **Figure 3B**). The ratios of exploration between unknown and now-known congener were similar in both groups, revealing no deleterious effect of social memory (*p* = 0.99) in CRE mice (63.4 ± 3 %) when compared to WT mice (63.5 ± 3%). Nevertheless, the total time of congener explorations was still found significantly lower (*p* < 0.0001) in CRE animals (83 ± 6 s) than in controls (139 ± 10 s, **Figure 3B**).

### 3.5. Home cage activity

**Data collected from the actimeter revealed different patterns of circadian activity according to mice genotype (Figure 4**). Total activity is expressed as the variable “All meters”, which represents the sum of all distances traveled within the beam break system (**Figure 4A**). The total mouse activity recorded during the dark phase was significantly higher (*p* = 0.032) in CRE mice (X = 125.4 ± 18.7 m) as compared to WT (X = 77.4 ± 10.8 m). Regarding the light cycle, CRE mice displaying higher activity levels (X = 26.1 ± 4.4 m) than controls (X = 16.6 ± 3.2 m), but the difference between the two strains did not reach statistical significance (*p* = 0.088). On a complete circadian cycle, CRE mice exhibited a significantly higher (*p* = 0.034) home cage activity (X = 161.7 ± 24.7 m) as compared to controls (X = 100.8 ± 12.9 m).

**Figure 4.** Effect of *Cre* genotype compared to WT on the behavior in home cage activity (CRE n=14, WT n=15). A) Measurement of all activities over a 24-h period covering an activity phase (dark phase) of 12 h and a resting phase (light phase) of 12 h. B) Displacement over the 24-h period. C) Fine movement (immobile activities like scratching and grooming) over the 24-h period. D) Sleeping percentage over the 24-h period. Data are shown as mean ± SEM. \**p* < 0.05, \*\**p* < 0.01 and \*\*\**p* < 0.001, significantly different from control (Fisher *post hoc* test)

As seen in **Figure 4B**, locomotor activity (“Ped meters”) counted for the major part of the “All meters” variable measured in **Figure 4A**, which reflected the total activity. The sum of direct locomotions recorded during the dark phase was found to be significantly higher (*p* = 0.043) in CRE mice (X = 120.3 ± 19.8 m) when compared to controls (X = 72.7 ± 11.2 m). The sum of direct locomotions performed during the light cycle revealed CRE mice displaying higher levels of locomotor activity (X = 21.3 ± 3.7 m) than controls (X = 13.6 ± 2.8 m) but the difference did not reach statistical significance (*p* = 0.115). Over the entire circadian cycle, CRE mice performed significantly more (*p* = 0.039) ambulatory activity (X = 141.5 ± 23.1 m) than WT (X = 86.4 ± 11.9 m).

The sum of all the displacements recorded due to fine activity was measured over the 24 h period (**Figure 4C**). During the dark phase, fine activity was found significantly higher (*p* = 0.039) in CRE mice (X = 15.3 ± 1.4 m) when compared to controls (X = 11.5 ± 1.1 m). Interestingly, CRE mice showed two peaks of immobile activity 3 h after the beginning of the dark phase and 2 h before the light phase (beginning of the sleeping phase). As in the dark phase, CRE mice exhibited more fine activity (X = 4.81 ± 0.7 m) than controls (X = 3 ± 0.4 m) during the light phase (*p* = 0.023). Another increase of immobile activity was found in CRE mice when compared to controls during the light phase at a specific time interval (from 8:00 P.M. to 9:00 P.M.). This phenomenon is similar to the peak observed during the dark phase which was also detected 2 to 3 h before the phase switch. Regarding the whole circadian cycle, CRE mice were found to perform significantly more (*p* = 0.022) fine activity (X = 20.1 ± 2 m) than controls (X = 14.4 ± 1.3 m) over a complete circadian cycle.

As reported in **Figure 4D**, the sleep sequences in CRE mice were greatly reduced during the dark phase when compared to those of control mice (*p* = 0.0005). It should be noted that two particular periods of arousal were concomitantly found to the two peaks of locomotor and fine activity previously highlighted during this dark phase, and thereby could contribute to the differences observed between the both strains. The sleep sequences in CRE mice were also reduced during the light phase when compared to those of control mice: the average sleep percentage was found at 95 ± 0.6 % in WT mice, while it was found at 91.1 ± 1 % in CRE mice (*p* = 0.0037). This was also the case for the whole circadian cycle: the average sleep percentage was found at 87.1 ± 1 % in WT mice, while it was found at 79.0 ± 1.7 % in CRE mice (*p* = 0.0002), which clearly shows different patterns of sleep during the whole circadian cycle between both CRE and WT.

### 3.5. Barnes maze

Mice spatial learning performances were assessed using a modified Barnes maze. Animals were given three trials a day for three consecutive days to learn the location of the escape hole. Learning performances and retention memory of both strains are shown in **Figure 5**. Concerning the acquisition period (Day 1 to Day 3, T1 to T9), the two-way ANOVA with repeated measures revealed at first a strong time effect (*p* < 0.0001), indicationg that both strains were able to learn the location of the escape hole as shown by reduction of time to perfrom the task. Nevertheless, looking to the second main factor, a clear genotype effect (*p* = 0.0049) was found, CRE mice spending significantly more time to find the escape than WT. Finally, no interaction between both factors was found (*p* = 0.23).

**Figure 5.**
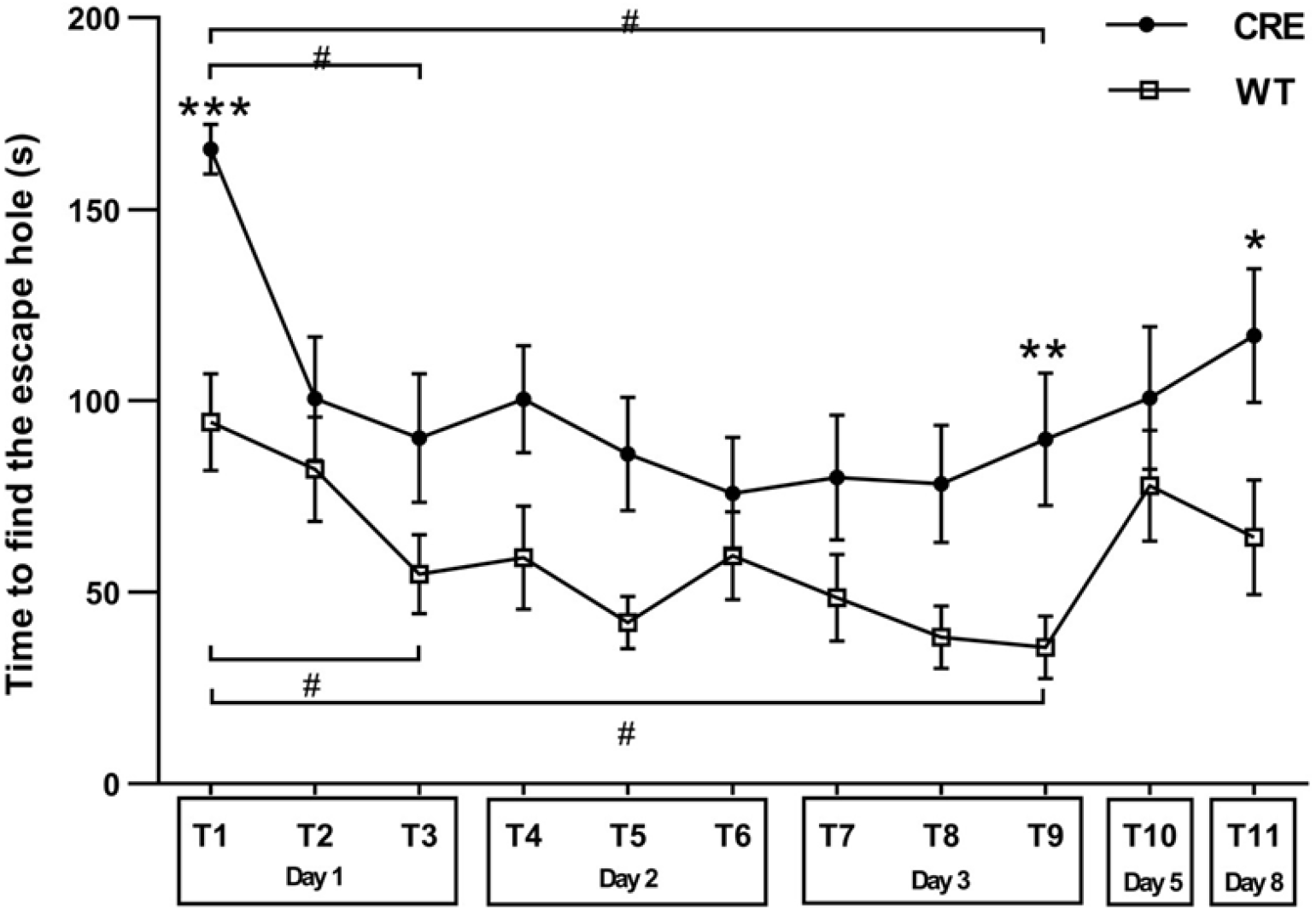
Effect of *Cre* genotype compared to WT on the latency to find the escape hole in the modified Barnes maze (CRE n=14, WT n=15). Data are reported as mean ± SEM. \**p* < 0.05, \*\**p* < 0.01, \*\*\**p* < 0.001, significantly different from control (unpaired *t* test post-hoc test). # p < 0.05 intra-group significant difference (Tuckey test)

For short-term learning, *post-hoc* Tuckey’s t-tests applied at 5% probability performed on trial 1 *vs*. trial 3 revealed a significant decrease in time to locate the escape hole in both groups. This indicated that short-term spatial learning took place since the first day of testing (mean difference of 39.7 with a critical difference of 37.2 for WT; mean difference of 75.4 with a critical difference of 49.3 for CRE; *p* < 0.05). *Post-hoc* Tuckey’s *t* tests also reached significance for both groups when comparing trial 1 *vs*. trial 9 (last trial of the acquisition period), suggesting continued learning over time (mean difference of 58.9 with a critical difference of 37.2 for WT; mean difference of 75.8 with a critical difference of 49.3 for CRE; *p* < 0.05). Thus, both groups evolved in time according to the same pattern. Both CRE and WT demonstrated short and long-term learning, but not at the same rate, with CRE mice exhibiting systematically longer durations to find the escape hole. Trial 1 has an intrinsic specificity, since none of the animals had ever been exposed to the maze before, which allowed to compare the performance of both groups at this particular time. Here, the unpaired *t* test showed a clear genotype effect (p < 0.0001), where CRE mice required up to 165 ± 6 s before finding the escape hole, as compared to a mean time of 94 ± 12 s for control mice to perform the same task. Finally, when looking for exploratory behavior in the last trial of the acquisition period (T9: third trial of the third day), a significant difference was still found. CRE mice required up to 90 ± 17 s when control mice showed a duration of 35 ± 8 s (*p* < 0.0073) to escape the maze’s aversive conditions. Two probe tests were performed on Day 5 and Day 8. No difference between both groups was found on Day 5 (*p* = 0.33). However, the Day 8 probe test revealed a deleterious effect of long-term memory in CRE mice when compared to controls (*p* = 0.03), since the time needed to find the escape hole was approximatively doubled in CRE mice (117 ± 18 s) in comparison with controls (64 ± 15 s).

## 4. Discussion

### 4.1. Hyperactivity in CRE mice

All performed tests pointed towards the hyperactive trait of CRE mice. This was especially noticeable for home cage activity where CRE mice walked longer distances than WT mice and had significantly more stationary activity including grooming and scratching than WT mice (**Figure 4**). In addition, CRE mice exhibited state anxiety behavior (**Table 2**) rather than trait anxiety behavior. The hyperactive trait of CRE mice most likely influenced their performance in all tests requiring attention and retention. CRE mice showed short-term memory problems (**Figure 1**), learning ability deficits and long-term memory problems (**Figure 5**). In addition, this disruption of information acquisition inevitably led to an impact on their performance in the object recognition test (**Figure 3**). Since the mice explored the objects significantly less as compared to WT, they had difficulties memorizing the familiar object and thus discriminating it from the newly introduced one.

CRE mice performed poorly in Barnes maze since their hyperactivity trait apparently blocked them from learning and retaining information (**Figure 5**). There is evidence in the literature that hyperactivity can interfere with escape latency, distance and speed in behavioral tests [15]. However, since CRE mice also performed badly in object recognition test, it is possible that they also have a visual deficit in addition to their hyperactive trait. Further tests that assess vision could be needed in order to rule out low vision ability in CRE mice. However, the retinal structure and organization of GLAST-CreERT2 mice appeared fully normal [4, 16], and routine morphological observation of the anterior chamber did not reveal corneal or lens opacification in CRE mice. In addition, Barnes maze can discriminate learning and retention abilities indepedently of visual abilities [17]. Thus, it is more likely that the hyperactive trait of CRE mice led to the low scores in the Barnes maze and object recognition tests, rather than global visual deficiency.

At the olfactory level, CRE mice were able to smell, as they performed well in buried cookie test (**Supplementary Figure S1**) and showed high interest in female urine odor in odor discrimination test (**Supplementary Figure S2**). They performed well in sociability and social novelty test (**Figure 3**), most probably due to a well-functioning olfactory system. However, they weren’t as interested in rose odor as in female urine and were unable to discriminate 1 % lemon additive (**Supplementary Figure S2**). This is most probably due to their hyperactivity trait rather than a deficient sensory entry. This was clearly visible in the habituation phase of the buried cookie test, when the cookie is not buried, but placed in a visible location in the cage. CRE mice took 4-times longer time to find the cookie compared to control. This was very likely not due to impaired sensory entry but rather inattention since the CRE mice were able to find the buried cookie. This hyperactive trait rather than olfactory deficits might also be linked to the lower interest for congeners identified in sociability tests (**Figure 3**).

### 4.2. Uncertainties about the mechanism of action

Noxious CRE effects have been documented over the past decade. First, CRE integration in mouse genome is random and can lead to multiple copies on single-chromosomal loci, as well as potentially disrupting endogenous gene expression [18]. Our CRE mice were generated using the bacterial artificial chromosome technology that are thought to minimize position effects on variegation and random recombination [4]. Even if we cannot fully rule out this possibility, it is unlikely that our CRE mice exhibit hyperactivity due to random gene disruption. *Cre* expression has been shown to induce DNA damage and apoptosis in the absence of genuine *loxP* sites [19, 20]. Toxic effects associated with *Cre* expression have been observed in various systems including the nervous system [21-23]. Moreover, TAM-inducible CRE expression has also been demonstrated to induce DNA damage in absence of valid *loxP* site [24]. However, the behavioral profile of our CRE mice is independent of the TAM treatment as both uninduced and induced CRE mice exhibited the same hyperactive trait (**Supplementary Figure S3**). This is in line with previous work in the field [25], where authors observed that short TAM treatments used to activate the CRE do not have a measurable impact on adult neurogenesis or on their performance in behavioral tests. In addition, in the original publication, no CRE recombinase activity was detected after vehicle injection (see Figure 3 in [4]), indicating absence of TAM-independent CreER^T2^ activity. Hyperactive trait is therefore neither linked to estrogen treatment, nor to detectable CRE recombinase activity that could have targeted invalid and degenerated *loxP* sites that are abundant in mouse genome [23].

The question whether accumulation of *cre* mRNA or inactive CRE recombinase protein could exert extra-genomic activity is still unanswered and the mechanism underlying the hyperactive behavior of our CRE mice is still to be deciphered. Interestingly, when we generated specific glia-specific conditional knockout mice based on this CRE recombinase system, we did not observe any hyperactive traits [7]. This would support the hypothesis of low CRE recombinase activity or that *cre* RNA accumulation might impair attention and activity in this CRE mice.

### 4.3. Importance of behavioral analysis of CRE transgenic mice

The hyperactivity trait observed in our CRE mice prevents it’s use as a negative control group in behavioral phenotyping of previously published conditional mutant mice [7]. In addition, the abnormal behavioral profile reported in this strain of CRE mice could indicate that CRE mice in general suffer from overlooked cognitive deficits. Indeed, there is little to no information on the specific behavior of CRE mice in the literature. In most behavioral studies, the group of interest (cKO) is usually compared to WT littermates, rather than CRE. Some of these studies showed that their group of interest was hyperactive when compared to *WT* controls [26-29]. Ignoring the behavioral status of CRE mice may generate a bias in the behavioral analysis of cKO mice by identifying a specific trait as a consequence of the targeted gene deletion while it might be linked in fact to the CRE recombinase activity. CRE group should therefore be analyzed in parallel to WT and cKO group in behavioral phenotyping experiments to avoid misinterpretation of gene deletion consequences. Moreover, similar results of specific tests might be due to different behavioral traits. For instance, CRE mice and previously characterized cKO mice [7] exhibited similar behavioral patterns in some of the tests, such as equal exploration time of objects and subjects, delay before departure in Barnes maze, and low proper alternation rates. However, these results observed were not caused by the same reason and were rather linked to hyperactivity and attention errors in CRE mice, as opposed to anxiety in the cKO mice that did not exhibit any hyperactive behavior [7]. Home cage activity and open field tests appear therefore as essential tests to easily discriminate between hyperactivity and anxiety and to avoid misinterpretation of behavioral profile linked to gene deletion consequences.

### 4.4. CRE mice as a model of attention-deficit hyperactivity disorder?

The hyperactive trait of CRE mice might serve as a model for attention deficit and hyperactivity disorder (ADHD). Here we showed that CRE mice were hyperactive, impulsive, and lacked motivation and attention, coinciding with ADHD which is generally characterized by inattention, hyperactivity and impulsivity, and deficits in motivation [30]. Under normal circumstances, the prefrontal cortex regulates attention, behavior, and emotion. ADHD is characterized by poor impulse control, weak sustained attention, and heightened distractibility, and has been linked to deficits in prefrontal cortex functioning [31]. In a stressed state, there is excessive dopamine and accompanied norepinephrine release which impairs prefrontal cortex abilities [32]. The pharmalogical treatment for ADHD is based on methylphenidate that blocks the activity of dopamine transporter and norepinephrine transporter, leading to increased availability of catecholamines in the synaptic cleft and to improved dopaminergic synapse functioning thereby restoring attention. Unlike amphetamine, methylphenidate does not induce dopamine release from presynaptic vesicles [33]. Animal models are critical to understand the pathological mechanisms involved in ADHD and to allow assessment and development of new therapeutics. The currently existing models are heterogeneous regarding physiopathological alterations, behavioral symptoms and response to medication [10, 34-36]. Indeed, no existing rodent model captures all aspects of ADHD. We demonstrated here that CRE mice exhibited impulsive and hyperactive traits. Their poor performances in attention-based tests were exacerbated when visual entry was required for a specific task (*i*.*e*., finding visible cookie in habituation phase). Those poor performances were linked to attention deficit rather than functional visual problems as they were able to oriente themselves based on clues in the Barne’s maze or discriminate illumination changes in the open field test. As we did not observe any other visual deficit in CRE mice, vision-based attention was therefore especially affected in CRE mice. In another study, authors identified ADHD-like behavior using vision-based attention tests in mutant mice exhibiting a duplicated retino-collicular map [37]. As those mice did not exhibit other visual impairement, it underlies the importance of superior colliculus organisation in vision-based attention. In addition, we pointed out a significant reduction of sleep periods for CRE mice (**Figure 4D**). Sleep disturbance is associated with ADHD in patients [38] and could also have affected CRE mice performances requiring attention and memorization. Interestingly, it was recently shown that sleep-deprived mice could serve as models of ADHD [39]. Altogether, CRE mice exhibit specific vision-based attention deficits associated with sleep disturbance, hyperactivity and impulsivity traits that affect their exploration, learning and retention abilities. CRE mice could therefore be proposed as an ADHD model in view of their behavioral response. Further experiments and studies would be needed to assess their adequation level with ADHD condition regarding physiopathological and pharmacological aspects to fullfull the list of criteria for an optimal animal model of ADHD as suggested previously [34].

## Author Contributions

Conceptualization, F.D., A.E.H., F.T.Y., M.C.L., T.C.; methodology, F.D., A.E.H., A.H., F.Dj. V.B., I.D.; validation, F.D., A.E.H., A.H., F.T.Y., M.C.L., T.C.; formal analysis, F.D., A.E.H., A.H., investigation, F.D., A.E.H., A.H.; ressources, F.Dj., T.O., C.M., F.T.Y., M.C.L., T.C.; data curation, F.D., A.E.H., A.H., F.T.Y., M.C.L., T.C.; writing—original draft preparation, F.D., A.E.H., A.H.; writing—review and editing, F.Dj., V.B., I.D., T.O., C.M., M.V., F.T.Y., M.C.L., F.T.Y., T.C.; visualization, F.D., A.E.H., A.H.; supervision, M.C.L., T.C.; project administration, F.T.Y, M.C.L., T.C.; funding acquisition, F.T.Y., T.C. All authors have read and agreed to the published version of the manuscript.

## Funding

This research was funded by the Région Lorraine and FEDER, Fondation Alzheimer, LUE IMPACT Biomolecules, and the French Ministry of Higher Education, Research and Innovation (MESRI). A.E.H.’s thesis fellowship was provided by the MESRI.

## Institutional Review Board Statement

The study was conducted according to the guidelines of the Declaration of Helsinki, and approved by the Ethics Committee of the University of Lorraine (protocol APAFIS #12079-201711081110404v2, February 02, 2018).

## Informed Consent Statement

Not applicable.

## Data Availability Statement

All data is contained in the article.

## Acknowledgments

The authors thank F. W. Pfrieger for providing Glast-CreERT2 mice.

## Conflicts of Interest

The authors declare no conflict of interest.

## Notes

### Competing Interest Statement

The authors have declared no competing interest.

## References

1. Sternberg, N.; Hamilton, D. Bacteriophage P1 site-specific recombination. I. Recombination between loxP sites. J. Mol. Biol. 1981, 150, 467–486.

2. Sauer, B.; Henderson, N. Site-specific DNA recombination in mammalian cells by the Cre recombinase of bacteriophage P1. Proc. Natl. Acad. Sci. U.S.A. 1988, 85, 5166–5170.

3. Tsien, J. Z.; Chen, D. F.; Gerber, D.; Tom, C.; Mercer, E. H.; Anderson, D. J.; Mayford, M.; Kandel, E. R.; Tonegawa, S. Subregion- and cell type-restricted gene knockout in mouse brain. Cell 1996, 87, 1317–1326.

4. Slezak, M.; Göritz, C.; Niemiec, A.; Frisén, J.; Chambon, P.; Metzger, D.; Pfrieger, F.W. Transgenic mice for conditional gene manipulation in astroglial cells. Glia 2007, 55, 1565-1576.

5. Indra, A.K.; Warot, X.; Brocard, J.; Bornert, J.M.; Xiao, J.H.; Chambon, P.; Metzger, D. Temporally-controlled site-specific mutagenesis in the basal layer of the epidermis: comparison of the recombinase activity of the tamoxifen-inducible Cre-ER(T) and Cre-ER(T2) recombinases. Nucleic Acids Res. 1999, 27, 4324–4327.

6. Li, M.; Snider, B.J. Chapter 1 - Gene Therapy Methods and Their Applications in Neurological Disorders. In Gene Therapy in Neurological Disorders; Li, M. & Snider, B.J. (eds); Academic Press, Cambridge, MA, U.S.A., 2018, pp. 3–39.

7. El Hajj, A.; Herzine, A.; Calcagno, G.; Désor, F.; Djelti, F.; Bombail, V.; Denis, I.; Oster, T.; Malaplate, C.; Vigier, M.; Kaminski, S.; Pauron, L.; Corbier, C.; Yen, F.T.; Lanhers, M.-C.; Claudepierre, T. Targeted suppression of lipoprotein receptor LSR in astrocytes leads to olfactory and memory deficits in mice. Int. J. Mol. Sci. 2022, 23, 2049.

8. Nigg, J.T. Attention-deficit/hyperactivity disorder and adverse health outcomes. Clin. psychol. rev. 2013, 33, 215–228.

9. Shaw, M.; Hodgkins, P.; Caci, H.; Young, S.; Kahle, J.; Woods, A. G.; Arnold, L.E. A systematic review and analysis of long-term outcomes in attention deficit hyperactivity disorder: effects of treatment and non-treatment. BMC Med. 2012, 10, 99.

10. Rahi, V.; Kumar, P. Animal models of attention-deficit hyperactivity disorder (ADHD). Int. J. Dev. Neurosci. 2021, 81, 107–124.

11. Leger, M.; Quiedeville, A.; Bouet, V.; Haelewyn, B.; Boulouard, M.; Schumann-Bard, P.; Freret, T. Object recognition test in mice. Nat. Protoc. 2013, 8, 2531–2537.

12. Kaidanovich-Beilin, O.; Lipina, T.; Vukobradovic, I.; Roder, J.; Woodgett, J.R. Assessment of social interaction behaviors. Journal of visualized experiments: JoVE 2011, 48, 2473.

13. Pack, A.I.; Galante, R.J.; Maislin, G.; Cater, J.; Metaxas, D.; Lu, S.; Zhang, L.; Von Smith, R.; Kay, T.; Lian, J.; Svenson, K.; Peters, L.L. Novel method for high-throughput phenotyping of sleep in mice. Physiol. Genom. 2007, 28, 232–238.

14. Youn, J.; Ellenbroek, B.A.; van Eck, I.; Roubos, S.; Verhage, M.; Stiedl, O. Finding the right motivation: genotype-dependent differences in effective reinforcements for spatial learning. Behav. Brain Res. 2012, 226, 397–403.

15. Baeta-Corral, R.; Giménez-Llort, L. Persistent hyperactivity and distinctive strategy features in the Morris water maze in 3xTg-AD mice at advanced stages of disease. Behav. Neurosci. 2015, 129, 129–137.

16. de Melo, J.; Miki, K.; Rattner, A.; Smallwood, P.; Zibetti, C.; Hirokawa, K.; Monuki, E. S.; Campochiaro, P. A.; Blackshaw, S. Injury-independent induction of reactive gliosis in retina by loss of function of the LIM homeodomain transcription factor Lhx2. Proc. Natl. Acad. Sci. U.S.A. 2012, 109, 4657–4662.

17. Yassine, N.; Lazaris, A.; Dorner-Ciossek, C.; Després, O.; Meyer, L.; Maitre, M.; Mensah-Nyagan, A.G.; Cassel, J.-C.; Mathis, C. Detecting spatial memory deficits beyond blindness in tg2576 Alzheimer mice. Neurobiol. Aging 2013, 34, 716–730.

18. Heaney, J.D.; Bronson, S.K. Artificial chromosome-based transgenes in the study of genome function. Mamm. Genome 2006, 17, 791–807.

19. Schmidt, E.E.; Taylor, D.S.; Prigge, J.R.; Barnett, S.; Capecchi, M.R. Illegitimate Cre-dependent chromosome rearrangements in transgenic mouse spermatids. Proc. Natl. Acad. Sci. U.S.A. 2000, 97, 13702–13707.

20. Janbandhu, V.; Moik, D.; Fässler, R. Cre recombinase induces DNA damage and tetraploidy in the absence of LoxP sites. Cell Cycle 2014, 13, 462–470.

21. Huh, W.J.; Mysorekar, I.U.; Mills, J.C. Inducible activation of Cre recombinase in adult mice causes gastric epithelial atrophy, metaplasia, and regenerative changes in the absence of “floxed” alleles. Am. J. Physiol.-Gastrointest. Liver Physiol. 2010, 299, G368–G380.

22. Harno, E.; Cottrell, E.C.; White, A. Metabolic pitfalls of CNS Cre-based technology. Cell Metab. 2013, 18, 21–28.

23. Pugach, E.K.; Richmond, P.A.; Azofeifa, J.G.; Dowell, R.D.; Leinwand, L. A. Prolonged Cre expression driven by the α-myosin heavy chain promoter can be cardiotoxic. J. Mol. Cell. Cardiol. 2015, 86, 54–61.

24. Lexow, J.; Poggioli, T.; Sarathchandra, P.; Santini, M.P.; Rosenthal, N. Cardiac fibrosis in mice expressing an inducible myocardial-specific Cre driver. Dis. Model Mech. 2013, 6, 1470–1476.

25. Rotheneichner, P.; Romanelli, P.; Bieler, L.; Pagitsch, S.; Zaunmair, P.; Kreutzer, C.; König, R.; Marschallinger, J.; Aigner, L.; Couillard-Després, S. Tamoxifen activation of Cre-recombinase has no persisting effects on adult neurogenesis or learning and anxiety. Front. Neurosci. 2017, 11, 27.

26. Ade, K.K.; Wan, Y.; Chen, M.; Gloss, B.; Calakos, N. An Improved BAC Transgenic fluorescent reporter line for sensitive and specific identification of striatonigral medium spiny neurons. Front. Syst. Neurosci. 2011, 5, 32.

27. Nelson, A.B.; Hang, G.B.; Grueter, B.A.; Pascoli, V.; Luscher, C.; Malenka, R.C.; Kreitzer, A.C. A comparison of striatal-dependent behaviors in wild-type and hemizygous Drd1a and Drd2 BAC transgenic mice. J. Neurosci. 2012, 32, 9119–9123.

28. Bodo, C.; Fernandes, C.; Krause, M. Brain specific Lamellipodin knockout results in hyperactivity and increased anxiety of mice. Sci. Rep. 2017, 7, 5365.

29. Bohuslavova, R.; Dodd, N.; Macova, I.; Chumak, T.; Horak, M.; Syka, J.; Fritzsch, B.; Pavlinkova, G. Pax2-Islet1 Transgenic Mice Are Hyperactive and Have Altered Cerebellar Foliation. Mol. Neurobiol. 2017, 54, 1352–1368.

30. Volkow, N.D.; Wang, G.-J.; Newcorn, J.H.; Kollins, S.H.; Wigal, T.L.; Telang, F.; Fowler, J.S.; Goldstein, R.Z.; Klein, N.; Logan, J.; Wong, C.; Swanson, J.M. Motivation Deficit in ADHD is associated with dysfunction of the dopamine reward pathway. Mol. Psychiatry 2011, 16, 1147–1154.

31. Arnsten, A.F.T. Catecholamine influences on dorsolateral prefrontal cortical networks. Biol. Psychiatry 2011, 69, e89–e99.

32. Wilens, T.E. Mechanism of action of agents used in attention-deficit/hyperactivity disorder. J. Clin. Psychiatry 2006, 67, 32–37.

33. Katzman, M.A.; Sternat, T. A Review of OROS methylphenidate (Concerta®) in the treatment of attention-deficit/hyperactivity disorder. CNS Drugs 2014, 28, 1005–1033.

34. Sagvolden, T.; Russell, V.A.; Aase, H.; Johansen, E.B.; Farshbaf, M. Rodent models of attention-deficit/hyperactivity disorder. Biol. Psychiatry 2005, 57, 1239–1247.

35. Sontag, T.A.; Tucha, O.; Walitza, S.; Lange, K.W. Animal models of attention deficit/hyperactivity disorder (ADHD): a critical review. Atten. Defic. Hyperact. Disord. 2010, 2, 1–20.

36. Russell, V.A. Overview of animal models of attention deficit hyperactivity disorder (ADHD). Curr. Prot. Neurosci. 2011, 9, 9–35.

37. Mathis, C.; Savier, E.; Bott, J.B.; Clesse, D.; Bevins, N.; Sage-Ciocca, D.; Geiger, K.; Gillet, A.; Laux-Biehlmann, A.; Goumon, Y.; Lacaud, A.; Lelièvre, V.; Kelche, C.: Cassel, J.C.; Pfrieger, F.W.; Reber, M. Defective response inhibition and collicular noradrenaline enrichment in mice with duplicated retinotopic map in the superior colliculus. Brain Struct. Funct. 2015, 220, 1573–1584.

38. Gruber, R. Sleep characteristics of children and adolescents with attention deficit-hyperactivity disorder. Child Adolesc. Psychiatr. Clin. N. Am., 2009, 18, 863–876.

39. Yaoita, F.; Namura, K.; Shibata, K.; Sugawara, S.; Tsuchiya, M.; Tadano, T.; Tan-No, K. Involvement of the hippocampal alpha2A-adrenoceptors in anxiety-related behaviors elicited by intermittent REM sleep deprivation-induced stress in mice. Biol. Pharm. Bull., 2020,

