## Supplementary figures for "CRE mice exhibit hyperactive and impulsive behavior affecting their learning and retention performances": 0 FIGURE S all.pptx

### Slide 1
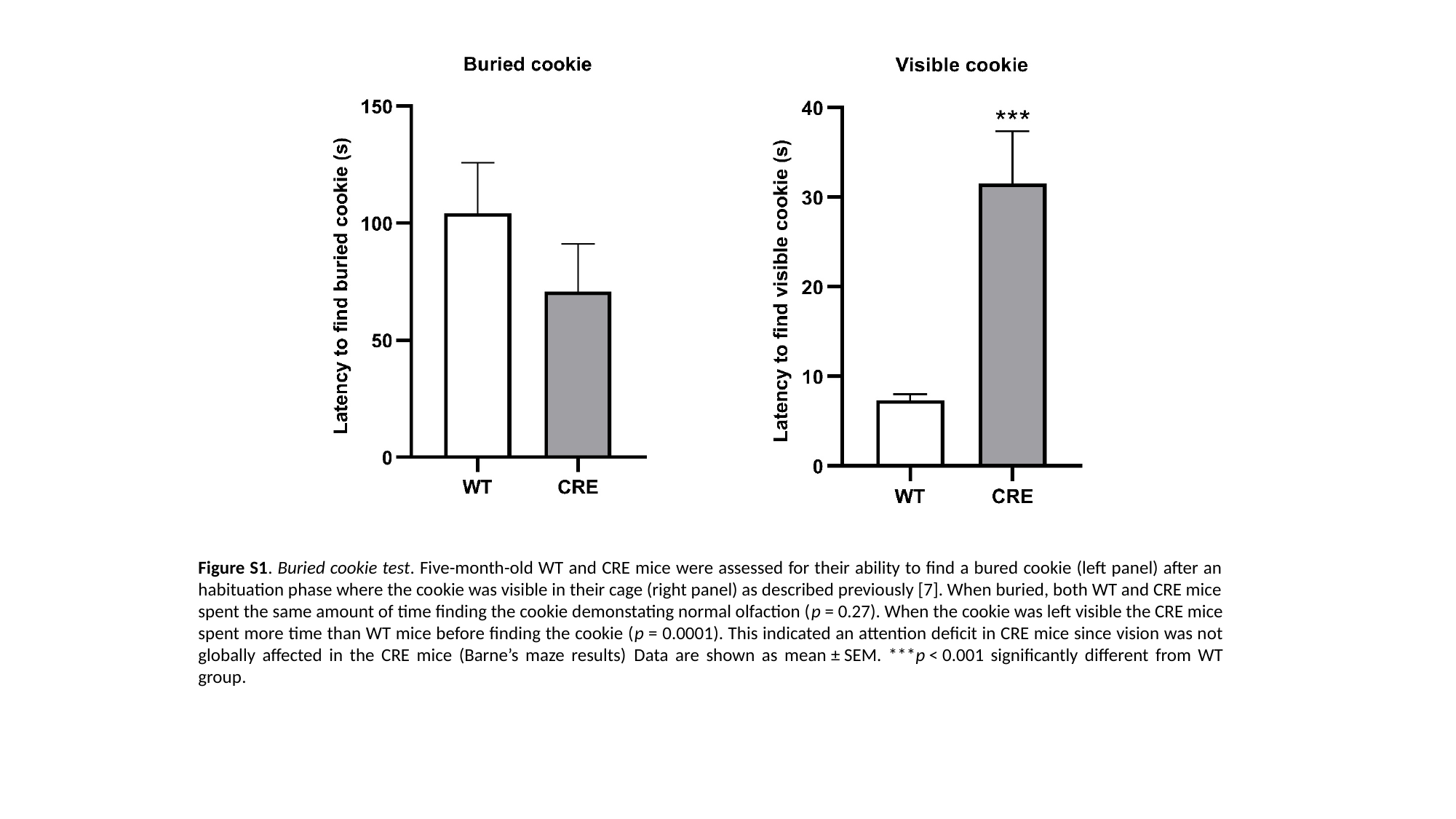

Figure S1. Buried cookie test. Five-month-old WT and CRE mice were assessed for their ability to find a bured cookie (left panel) after an habituation phase where the cookie was visible in their cage (right panel) as described previously [7]. When buried, both WT and CRE mice spent the same amount of time finding the cookie demonstating normal olfaction (p = 0.27). When the cookie was left visible the CRE mice spent more time than WT mice before finding the cookie (p = 0.0001). This indicated an attention deficit in CRE mice since vision was not globally affected in the CRE mice (Barne’s maze results) Data are shown as mean ± SEM. ***p < 0.001 significantly different from WT group.

### Slide 2
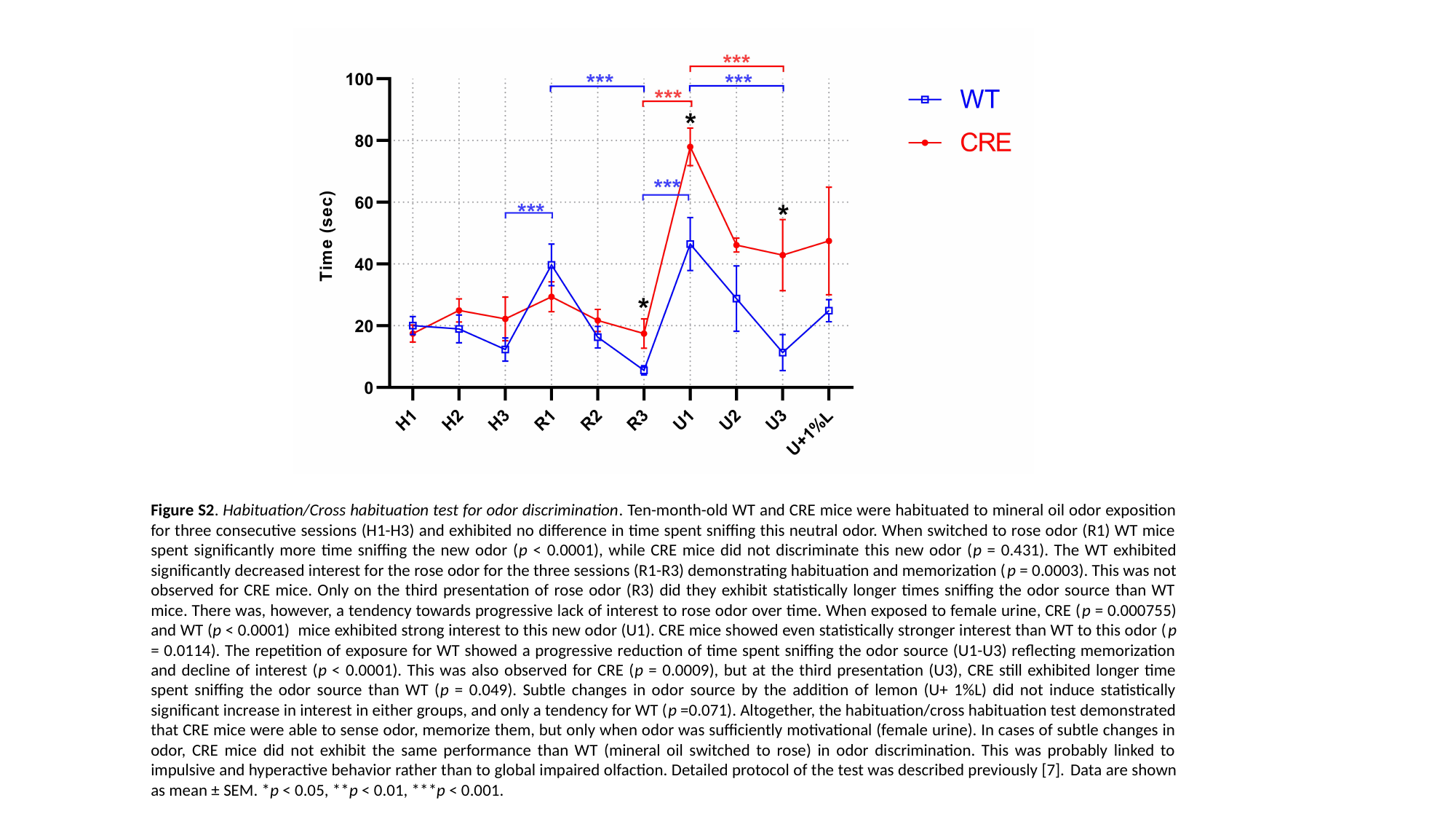

Figure S2. Habituation/Cross habituation test for odor discrimination. Ten-month-old WT and CRE mice were habituated to mineral oil odor exposition for three consecutive sessions (H1-H3) and exhibited no difference in time spent sniffing this neutral odor. When switched to rose odor (R1) WT mice spent significantly more time sniffing the new odor (p < 0.0001), while CRE mice did not discriminate this new odor (p = 0.431). The WT exhibited significantly decreased interest for the rose odor for the three sessions (R1-R3) demonstrating habituation and memorization (p = 0.0003). This was not observed for CRE mice. Only on the third presentation of rose odor (R3) did they exhibit statistically longer times sniffing the odor source than WT mice. There was, however, a tendency towards progressive lack of interest to rose odor over time. When exposed to female urine, CRE (p = 0.000755) and WT (p < 0.0001) mice exhibited strong interest to this new odor (U1). CRE mice showed even statistically stronger interest than WT to this odor (p = 0.0114). The repetition of exposure for WT showed a progressive reduction of time spent sniffing the odor source (U1-U3) reflecting memorization and decline of interest (p < 0.0001). This was also observed for CRE (p = 0.0009), but at the third presentation (U3), CRE still exhibited longer time spent sniffing the odor source than WT (p = 0.049). Subtle changes in odor source by the addition of lemon (U+ 1%L) did not induce statistically significant increase in interest in either groups, and only a tendency for WT (p =0.071). Altogether, the habituation/cross habituation test demonstrated that CRE mice were able to sense odor, memorize them, but only when odor was sufficiently motivational (female urine). In cases of subtle changes in odor, CRE mice did not exhibit the same performance than WT (mineral oil switched to rose) in odor discrimination. This was probably linked to impulsive and hyperactive behavior rather than to global impaired olfaction. Detailed protocol of the test was described previously [7]. Data are shown as mean ± SEM. *p < 0.05, **p < 0.01, ***p < 0.001.

### Slide 3
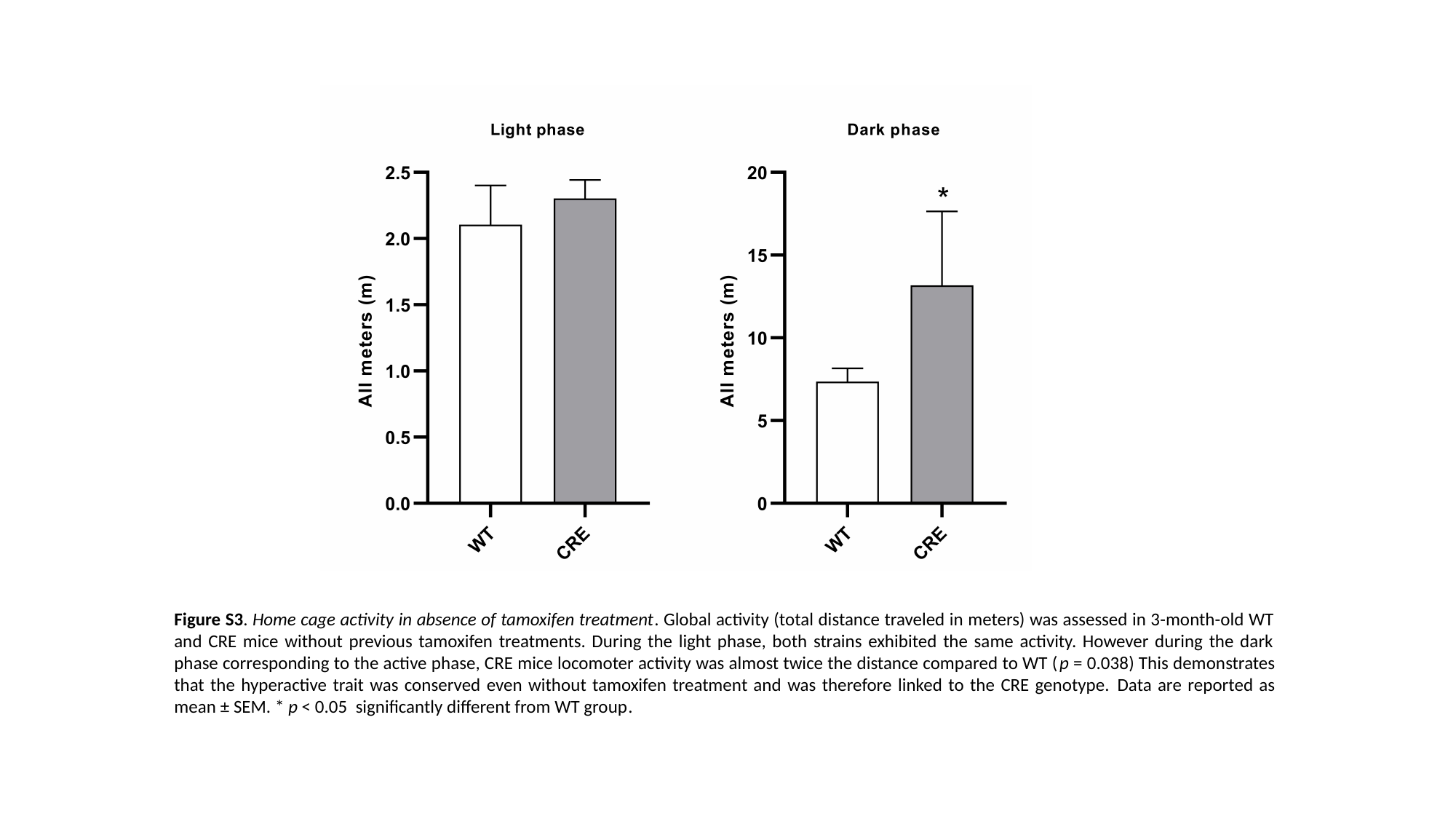

Figure S3. Home cage activity in absence of tamoxifen treatment. Global activity (total distance traveled in meters) was assessed in 3-month-old WT and CRE mice without previous tamoxifen treatments. During the light phase, both strains exhibited the same activity. However during the dark phase corresponding to the active phase, CRE mice locomoter activity was almost twice the distance compared to WT (p = 0.038) This demonstrates that the hyperactive trait was conserved even without tamoxifen treatment and was therefore linked to the CRE genotype. Data are reported as mean ± SEM. * p < 0.05 significantly different from WT group.
